# Microbiomes of North American Triatominae: the grounds for Chagas disease epidemiology

**DOI:** 10.1101/269845

**Authors:** Sonia M. Rodríguez-Ruano, Veronika Škochová, Ryan O. M. Rego, Justin O. Schmidt, Walter Roachell, Václav Hypša, Eva Nováková

## Abstract

Insect microbiomes influence many fundamental host traits, including functions of practical significance such as their capacity as vectors to transmit parasites and pathogens. The knowledge on the diversity and development of the gut microbiomes in various blood feeding insects is thus crucial not only for theoretical purposes, but also for the development of better disease control strategies. In Triatominae (Heteroptera: Reduviidae), the blood feeding vectors of Chagas disease in South America and parts of North America, the investigation of the microbiomes is in its infancy. The few studies done on microbiomes of South American Triatominae species indicate a relatively low taxonomic diversity and a high host specificity. We designed a comparative survey to serve several purposes: I) to obtain a better insight into the overall microbiome diversity in different species, II) to check the long term stability of the interspecific differences, III) to describe the ontogenetic changes of the microbiome, and IV) to determine the potential correlation between microbiome composition and presence in the vector gut of *Trypanosoma cruzi*, the causative agent of Chagas disease. Using 16S amplicons of two abundant species from the southern US, and four laboratory reared colonies, we showed that the microbiome composition is determined by host species, rather than locality or environment. The OTUs (Operational Taxonomic Units) determination confirms a low microbiome diversity, with 12-17 main OTUs detected in wild populations of *T. sanguisuga* and *T. protracta*. Among the dominant bacterial taxa are *Acinetobacter* and *Proteiniphilum* but also the symbiotic bacterium *Arsenophonus triatominarum*, previously believed to only live intracellularly. The possibility of ontogenic microbiome changes was evaluated in all six developmental stages and faeces of the laboratory reared model *Rhodnius prolixus*. We detected considerable changes along the host’s ontogeny, including clear trends in the abundance variation of the three dominant bacteria, namely *Enterococcus*, *Acinetobacter* and *Arsenophonus*. Finally, we screened the samples for the presence of *T. cruzi*. Comparing the parasite presence with the microbiome composition, we assessed the possible significance of the latter in the epidemiology of the disease. Particularly, we found a trend towards more diverse microbiomes in *T. cruzi* positive *T. protracta* specimens.

## Introduction

Insect microbiomes are recognized as a major factor determining host fitness and various phenotypic traits, including vectorial capacity in blood feeding species (Klepzig et al., 2009; Minard et al., 2013; Oliver and Martinez, 2014). While vector control strategies are now naturally venturing into the microbiome concept (Crotti et al., 2012; Saldaña et al., 2017), for some insect vectors and the allied pathogens the research focus has not yet fully passed the borders of epidemiological, surveillance and medical studies. One such example are kissing bugs of the subfamily Triatominae (Hemiptera: Reduviidae) and *Trypanosoma cruzi*, the protozoan parasite responsible for Chagas disease.

Triatominae are hemimetabolous insects with five nymphal instars that feed on vertebrate blood. There are over 130 extant Triatominae species with different biology, vectorial capacity and distribution, most of them found in South and Central America (Galvão et al., 2003). Those areas are the hot spots for Chagas disease, with nearly six million people affected (WHO, 2015). Among the seven main vector species of *T. cruzi* in Latin America (Hernández et al., 2016), the most attention has traditionally been paid to Rhodnius *prolixus* colonizing domestic habitats (Gourbière et al., 2012). For many decades, *R. prolixus* has also served as a model organism to study insect physiology, immunity, metabolism, and development (Nunes-da-Fonseca et al., 2017). Such a status has promoted a high availability of information on this Triatominae species, including a whole genome sequence (Mesquita et al., 2015), and has contributed to unraveling its role as *T. cruzi* vector.

The southern part of the United States is also endemic for this trypanosomiasis (Montgomery et al., 2014) and Chagas disease has recently been listed among the emerging diseases in this region (Edwards et al., 2017). Eleven Triatominae species are considered native to North America (Bern et al., 2011). The five species most frequently found are *Triatoma gerstaeckeri, T. indictiva, T. lecticularia, T. sanguisuga, and T. rubida* (Curtis-Robles et al., 2017a). So far, a modest number of both human and canine *T. cruzi* infections has been documented in Texas (Curtis-Robles et al., 2017a; Wozniak et al., 2015). The latest surveillance data, however, reveal the presence of Triatominae adults and nymphs in domestic and peridomestic areas, which points at a potential increase of human infections (Curtis-Robles et al., 2017a). In fact, current studies are already describing a growing number of human subjects locally infected with *T. cruzi* in southern US (Garcia et al., 2017; Gunter et al., 2017).

Since *T. cruzi* undergoes development exclusively in the intestinal tract of Triatominae vectors, it is exposed to interactions with gut microbes. Many studies to date have revealed an effect (either positive or negative) of microbiomes on the vectorial capacity of their insect hosts through different mechanisms such as resource competition, immunity modulation or antiparasitic secondary metabolite production (Cirimotich et al., 2011; Hegde et al., 2015; Jupatanakul et al., 2014). In Triatominae, the microbiomes interact with *T. cruzi* by direct contact, including competition for resources (Garcia et al., 2010), or indirectly by increasing the expression of antiparasitic molecules and humoral immune defense factors of the host (Azambuja et al., 2005; Weiss and Aksoy, 2011). While there are two major threads to commonly used chemical control of Triatominae vectors, i.e. developing resistance and recurrent infestations by sylvatic species, symbiont manipulation has been proposed as a sustainable alternative (Gourbière et al., 2012). For instance, trypanolitic activity of some Triatominae associated *Serratia marcescens* strains (Azambuja et al., 2004) and gene silencing using transformed *Rhodococcus rhodnii* have been explored (Taracena et al., 2015). However, any advancement in microbiome-based vector control is hampered by the lack of a solid background on Triatominae symbiosis arising from the basic knowledge on the microbiome characteristics.

Until now, microbiome data and their basic characteristics have been available for a limited number of South American Triatominae species from wild and laboratory colonies (da Mota et al., 2012; Díaz et al., 2016; Gumiel et al., 2015). Based on these data, Triatominae possess low-complexity microbiomes, with one or few dominant bacterial genera, which appear to be specific to certain hosts (da Mota et al., 2012; Díaz et al., 2016). The complexity of lab-reared insect microbiomes is even lower, but maintains most of the bacterial groups found in their wild counterparts (da Mota et al., 2012). While for other insects the microbiomes are known to vary along the host ontogeny (Duguma et al., 2015; Gimonneau et al., 2014; Sudakaran et al., 2012), the development of Triatominae microbiomes is completely unknown. Considering the ability of all the instars to transmit *T. cruzi* (Wozniak et al., 2015), possible changes in the microbiome composition along their ontogeny may be a key point towards the understanding of the disease transmission and its biological control. In this study, we extend the analysis of microbiome diversity and specificity to North American species from wild and laboratory colonies. We also take advantage of the model species *R. prolixus* and describe the microbiome variation in its different developmental stages. Finally, in order to establish the grounds for microbiome-based control strategy, we present the first insight into the microbiome - *T. cruzi* interface from wild populations of two broadly found North American species, *T. protracta* and *T. sanguisuga*.

## Materials and Methods

### The design of the sample set: taxonomy and origin

We used samples of four Triatominae species reared in long term laboratory colonies (*R. prolixus*, *T. vitticeps*, *T. protracta* and *T. recurva*) and field collected samples of two species, *T. protracta* and T. *sanguisuga*. Due to this arrangement we could address the main hypothesis (i.e. the host specific microbiome composition) in two different ways. First, we compared field sampled individuals of two species from different localities to see whether the microbiome composition is determined by the host taxonomy or rather the geography. Second, we compared microbiomes of four species from laboratory colonies to determine whether the significant interspecific differences can be detected after a long term cohabitation in the same environment. In addition to this comparative analyses, we used the *R. prolixus* colony to directly characterize the development of the microbial communites throughout the host ontogeny, i.e. across different developmental stages.

### Rearing conditions and microbiome ontogreny

The microbiome ontogeny was assessed as follows. Eggs, all five nymphal instars (L1-L5), and adults (L6) of *R. prolixus* were analyzed in triplicates as described hereafter. Upon faeces availability, filter paper collected faeces were processed in five replicates. The *R. prolixus* colony, established in 2010, originates from Brazil and has since been kept at the Faculty of Science, University of South Bohemia, Czech Republic. The colony is reared in plastic containers with mesh lids at a constant temperature of 27 °C and 70 % humidity under 16:8 hour light/dark cycles. The blood meal, commercially purchased defibrinated sheep blood supplemented with egg yolk (Núñez and Lazzari, 1990), is supplied in ten day intervals using artificial membrane feeding *ad libitum*.

### Interspecific diversity and stability of the microbioemes

The variability and host specificity of the microbial communities (i.e. the assumption that the specific microbiome profile is determined by the host species), was tested using laboratory reared colonies. In order to prevent downsizing of the colonies, only first and second instars were sampled in five replicates from four Triatominae species. The origins of the samples were as follows (see Table 1). The *R. prolixus* colony and *T. vitticeps* individuals were acquired from the above-described facility and were kept under the same breeding conditions. The colony of *T. vitticeps* originates from field-captured individuals sampled in Brazil and kept since 1995. The two other species, *T. protracta* and *T. recurva*, representing the North American (NA) Triatominae, were provided by the Southwestern Biological Institute, Tucson, AZ, USA. All the NA colonies were established by Justin Schmidt in 2010. *T. protracta* originated from Tucson, AZ, while *T. recurva* was sampled in Bisbee, AZ. The colonies are kept under simulated natural environmental conditions experienced by the insects in their native habitat. The temperature in the insectary is allowed to fluctuate with the weather, except with heating to maintain at least 4°C and cooling to maintain a temperature no higher than 32°C. Humidity is not being modified except during cooling conditions when maintained at 30%. Lighting is natural as provided by numerous windows of the insectary. Meals of rodent blood are provided every 14 days *ad libitum* as approved by the institutional review board of the Southwestern Biological Institute.

**Table 1.**
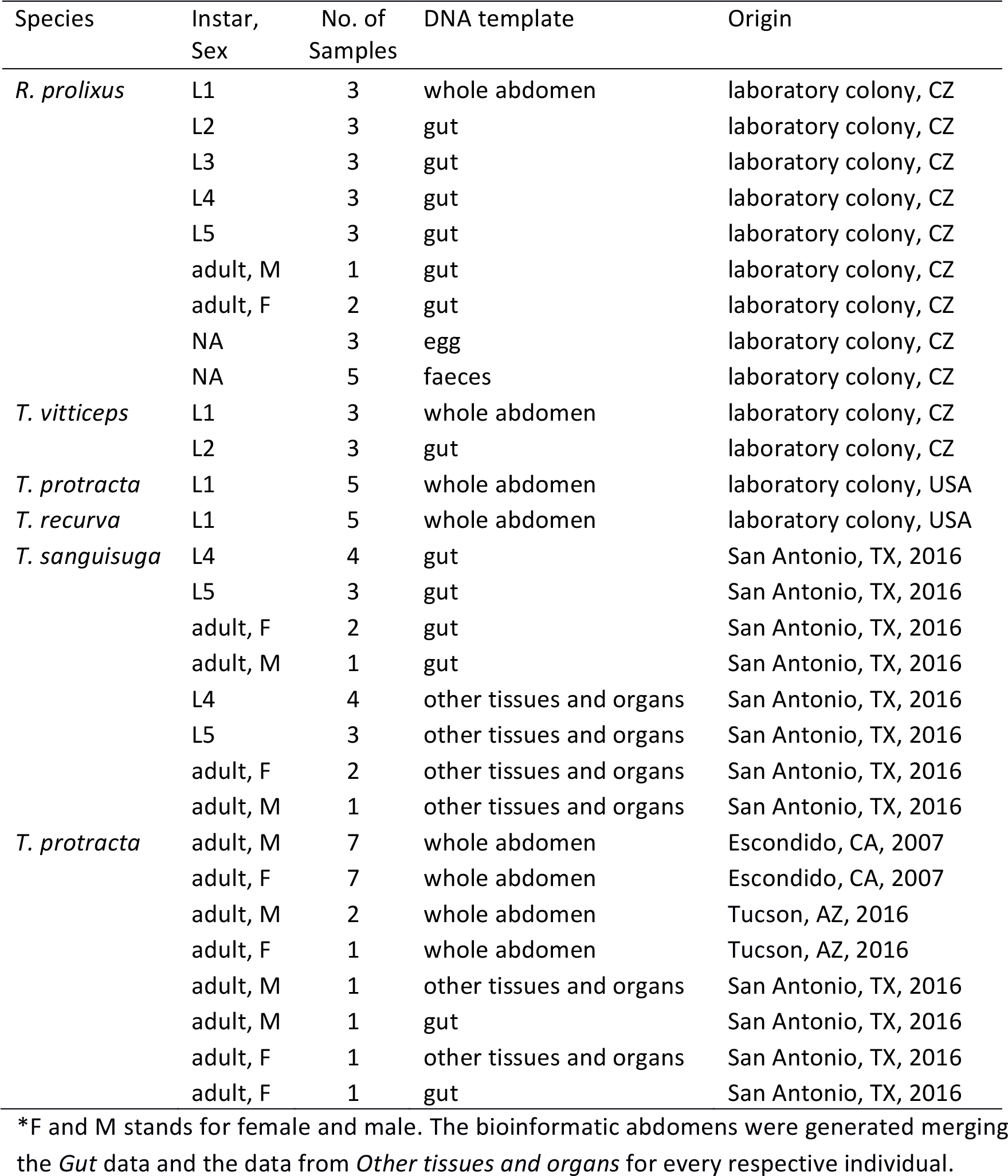
List of all the DNA isolates for which the 16S rRNA gene amplicon data were 533 generated in this study.

In addition to the laboratory-reared samples, we analyzed field collected samples to gain an initial insight into the diversity and ecology of the microbiome associated with the most widespread NA species. Triatominae sampled in the three most affected states, i.e. California, Arizona and Texas, were provided by Christiane Weirauch (UC Riverside, California) and Walter D. Roachell (US Army Public Health Command Central, Texas). The samples were morphologically assigned to two species: *T. sanguisuga* and *T. protracta*, and the determination was further confirmed by molecular data. The complete information on laboratory bred and field collected individuals used in this study is provided in Table 1.

### *Template extraction, molecular taxonomy p and* T. cruzi *diagnostics*

Colony individuals were sampled 4 days after feeding. Due to their size, the first instars were not dissected, and whole surface sterilized abdomens were used (see below for sterilization details). For the second and older instars in the ontogenetic survey, gut samples were acquired using a simple dissection protocol. The bugs were surface sterilized in ethanol followed by a rinse in sterile deionizied water and dissected in sterile phosphate buffer saline (PBS). The ventral cuticule was removed and the entire length of the gut was collected into individual tubes for each DNA extraction. The dissection ensured acquiring only bacteria from the gut, where the main interactions with the host and possible parasites are expected to occur, and it also reduced any external contamination.

For the field sampled individuals, we optimized the extraction protocol so that comparable microbiome profiles could be generated from ethanol preserved abdomens (provided by some of our collaborators and available in numerous collections worldwide) and fresh dissected samples. Two separate templates were used from the dissected samples, i.e. the gut and the other abdominal tissues and organs (Table 1). This protocol, followed by merging the corresponding microbiome data into a “bioinformatic abdomen” (see Data processing and statistical analyses), ensures the data comparability and provides gDNA gut specific templates for any future applications (i.e. qPCR and metagenomics). Total DNA was extracted from all the above described templates (Table 1) using the DNeasy Blood and Tissue (Qiagen, Hilden, Germany).

In order to verify the taxonomical determination of the 29 field collected individuals, we sequenced fragments of the COI gene amplified with the forward primers COIL6625 and/or COIL1490 and the reverse primer COIH7005 (Hafner et al., 1994). The reference sequence for *T. sanguisuga* was obtained from the adult individuals provided by Walter Roachel. The reference for *T. protracta* was sequenced from the laboratory colony (see above). A Maximum Likelihood (ML) analysis was performed with PhyML plugin in Geneious (Guindon et al., 2010; Kearse et al., 2012) to confirm the taxonomical assignment.

The field collected individuals were further screened for the presence of *T. cruzi*. The diversity of *T. cruzi* discrete typing units (DTUs), associated with geography, vector species, transmission and clinical outcomes (Hernandez 2016), spans five out of the six designated DTUs within United States (Garcia 2017). Three sets of diagnostic primers were thus employed in this study: one generally amplifying all *T. cruzi* DTUs (TCZ1/TCZ2; Moser et al., 1989) and the two other distinguishing for DTUI and the DTUII-VI group (ME/TC1 or TC2; Gumiel et al., 2015).

### Library preparation and microbiome sequencing

To generate data suitable for microbiome profiling, gDNA of 83 samples was amplified according to the EMP protocol (http://www.earthmicrobiome.org/protocols-and-standards/16s/), producing approximately 250 pb of the 16S rDNA V4 hypervariable region. Two negative controls of the extraction procedure and one blank control for PCR amplification were included in the sequencing. The libraries were then sequenced in a single run of Illumina MiSeq using the v2 chemistry (2×150 bp output mode).

### Data processing and statistical analyses

Forward and reverse paired-end reads were merged using USEARCH v7.0.1001 (fastq_mergepairs with fastq_minovlen set to 20; Edgar, 2013). Then the sequences were demultiplexed and their quality was checked using QIIME 1.8 (split_libraries_fastq.py with phred_quality_threshold set to 19; Caporaso et al., 2010b). The retained high quality sequences were aligned using the QIIME implementation of Pynast (Caporaso et al., 2010a) and trimmed to a final length of 251 bp using USEARCH (Edgar, 2013). Finally, the dataset was clustered at 100% identity to get a representative set for *de novo* OTU picking using the USEARCH global alignment option at 97 % identity (Edgar, 2013). Each OTU was assigned to different taxonomic levels using the BLAST algorithm (Camacho et al., 2009) against the SILVA 123 database (Quast et al., 2013). The obtained OTU table was filtered to remove singletons and very low abundant OTUs using QIIME (following the recommendations of Bokulich et al., 2013), and all non-bacterial, chloroplast and mitochondrial OTUs were filtered out. After this final filtering step, the OTU table was rarefied at 700 sequences per sample to normalize the dataset (as recommended in Weiss et al., 2017). The unrarefied data obtained for guts and other internal organs/tissues from each of the dissected wild sampled individuals were merged into “bioinformatic abdomens” using the QIIME script *collapse_samples.py* and finally normalized at 1000 sequences per sample.

All statistical tests were performed in R environment (R Development Core Team, 2014), mainly using functions from the vegan package (Oksanen et al., 2013). The overall similarity among microbiomes was analyzed based on Bray-Curtis dissimilarities and visualized in two dimensional space using Non-Metric Dimensional Scaling (NMDS; Minchin, 1987). Using the vegan Adonis function, we tested the statistical significance of the differences among microbiomes from distinct host instars and host species. For *T. protracta*, differences between the distribution of diversity indices (Shannon index and richness) of males and females and between *T. cruzi* positive and negative individuals were statistically evaluated using Kruskal-Wallis rank test (Kruskal and Wallis, 1952). All graphical outputs were generated using ggplot2 (Wickham, 2009).

## Results

### Sanger and Illumina data

For molecular determination, we obtained 25 fragments 381 bp long and 4 fragments 652 bp long of the COI sequences. ML analysis of these fragments (not shown) determined 10 individuals as *T. sanguisuga* and 19 as *T. protracta*. The representative sequences for each species were deposited in GenBank under the following accession numbers: XZ12345-6.

Illumina data were obtained for 42 samples from the laboratory reared colonies and 29 wild sampled individuals and 3 negative controls. The mean number of high quality merged reads passing the QC was 9279 (SD = 554). The extraction procedure controls resulted in extremely low read numbers (988 and 666). The first extraction control was dominated by OTU161 (*Segetibacter*) that did not occur in any of the Triatominae microbiome data. The second extraction control sample as well as the blank PCR sample (253 reads) were composed of a mixture of low abundant OTUs that were excluded from further analysis. The raw data are available under the study no. PRJEB25175.

### Microbiome development throughout the host ontogeny

The microbiome of *R. prolixus* undergoes major compositional changes along the host
ontogeny (Figures 1 and 2). Significant differences were found when comparing the microbiome of eggs, first four instars, fifth instar and adults, and faeces of this species (R^2^ = 0.61335, significance level of 0.001, Figure 1B). When compared pairwise, all the microbiome pairs bore significant differences (four pairs at 0.01 level of significance and two at 0.05, see Supplementary Data: Table 1). On average, the egg replicates showed the highest richness: 37 OTUs represented by numerous aerobes and facultative anaerobes. Two taxa equally dominated the microbial composition, namely *Enterococcus* and *Bartonella* (each accounting for 21.8 %). The third well represented taxon was *Methylobacterium* (18.2 %).

**Figure 1.**
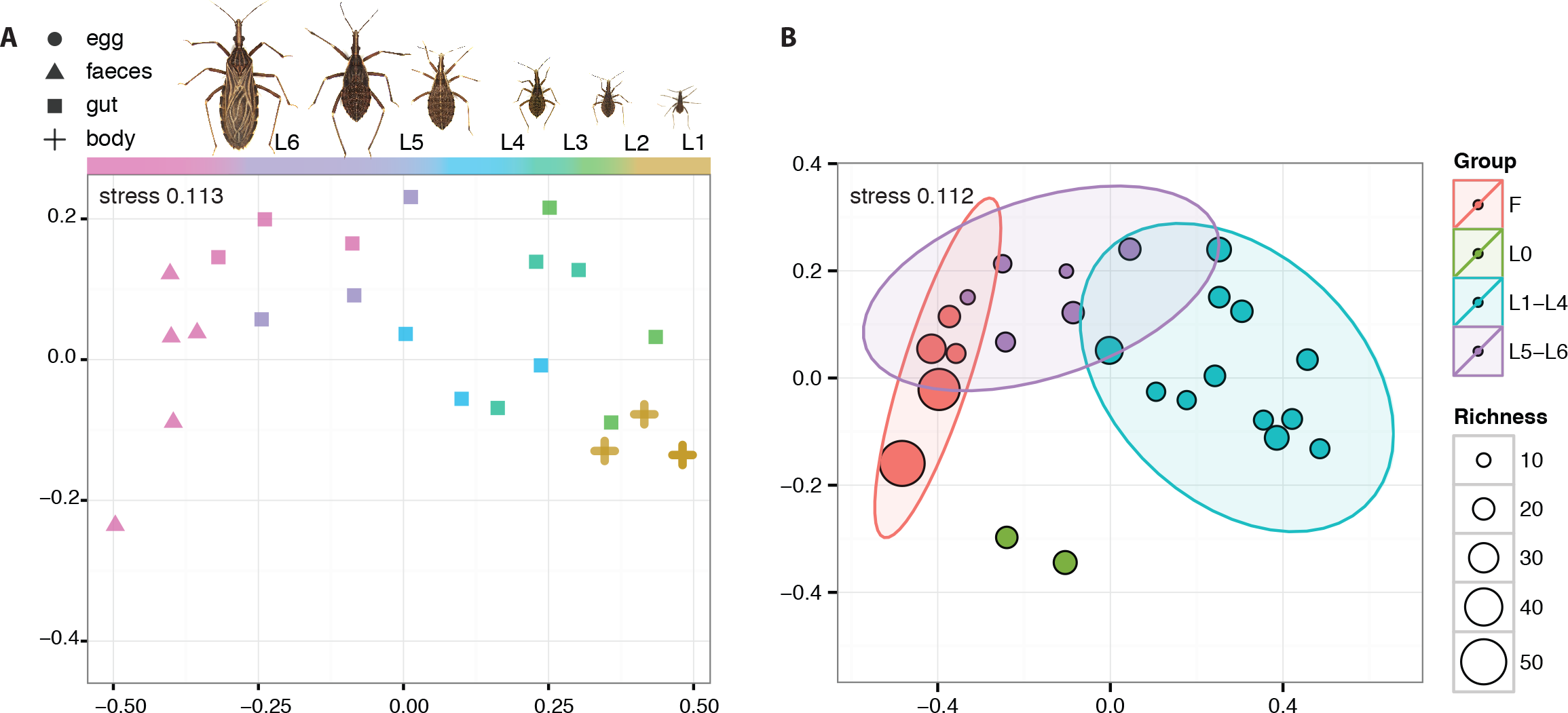
NMDS visualization of the ontogenetic shift of *R. prolixus* microbiome. F stands stands for faeces, L0 for eggs, L1-L5 for nymphal instars, L6 for adults.

Starting with the first nymphal instar the composition of the bacterial community changes dramatically (Figure 2). The richness of the microbiome drops in the nymphal stages to 13-19 OTUs (Figure 2B), and in the faeces it reaches 92 OTUs. During the ontogeny from L1 to an adult, the composition of the microbiome shows several consistent trends. The most pronounced of these trends are expressed in the quantities of the three dominant OTUs, *Enterococcus*, *Acinetobacter* and *Arsenophonus* (Figure 2B). While the proportion of *Enterococcus*/*Acinetobacter* in the microbiome steadily increases/decreases during the host ontogeny, the number of *Arsenophonus* reaches its peaks in L3 and then starts decreasing (Figure 2). The only dominant OTU without clear consistent pattern was *Macrococcus* which varied in numbers among the nymphal stages and was absent in adults.

**Figure 2.**
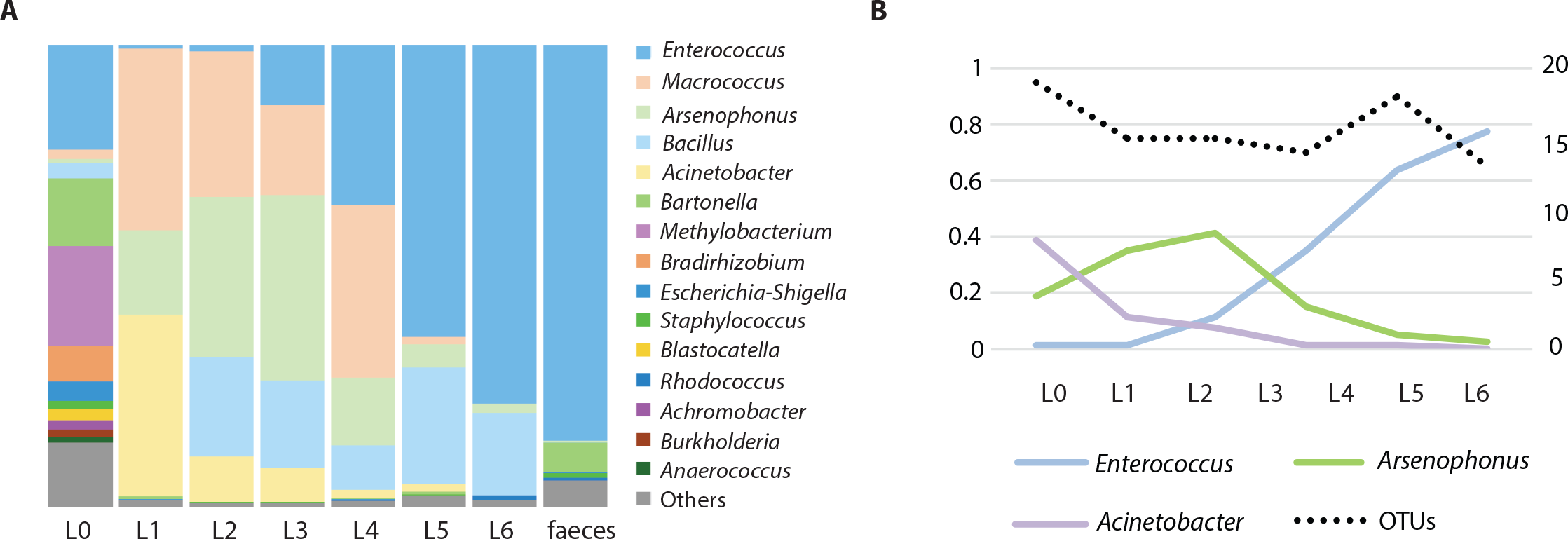
Microbiome of *R. prolixus* along its ontogeny. (A) Relative abundance of the main genera in different developmental stages. L0 stands for eggs, L1-L5 are nymphal instars, L6 represents adults. (B) Variation of the dominant OTUs; percentage of the total reads number (left axis) and the number of OTUs in individual stages (right axis).

### *Microbiomes of* T. protracta *and* T. sanguisuga

The two species from wild populations showed major differences between their microbiome profiles at the significance level of 0.001 (R^2^ = 0.289, Figure 3). Since the samples of *T. protracta* were mostly alcohol preserved, the two species were compared based on the microbial data retrieved from the entire abdomens and bioinformatically merged abdomen data. *T. sanguisuga* microbiomes retrieved from individuals of late nymphs (L4 and L5) and adults were composed of 17 OTUs with an averaged relative abundance higher than 1% (Supplementary Data: Table 2). Among these, *Arsenophonus* and *Acinetobacter* bacteria clearly dominated. On average the relative abundance of *Arsenophonus* in all analyzed *T. sanguisuga* individuals was 39.5%, reaching 82.5% in the L4 nymphs, while for *Acinetobacter* the average was 14.3% with the highest relative abundance value of 34.1% in the last nymphal instar L5. Four other taxa exhibited an average relative abundance over 3%, i.e. *Stenotrophomonas* (6.2%), *Rhodopseudomonas* (4.6%), *Pseudomonas* (4.4%) and *Sphingobacterium* (3.4%) (Supplementary Data: Table 2). *T. protracta* microbiome of 16 adult individuals was composed of 12 OTUs with an average relative abundance higher than 1% and the dominating OTU assigned to the genus *Proteiniphilum* (40.7%). Five OTUs showed an average relative abundance above 3%, i.e. *Dietzia* (11.6%), *Salmonella* (10.4%), *Peptoniphilus* (9.0%), Neisseriaceae (6.1%), *Mycobacterium* (3.7%). *Dietzia* was however recovered from a single individual of *T. protracta* reaching the abundance of 97% of all reads generated from this particular specimen (removed as an outlier from the *T. cruzi* related analysis).

**Figure 3.**
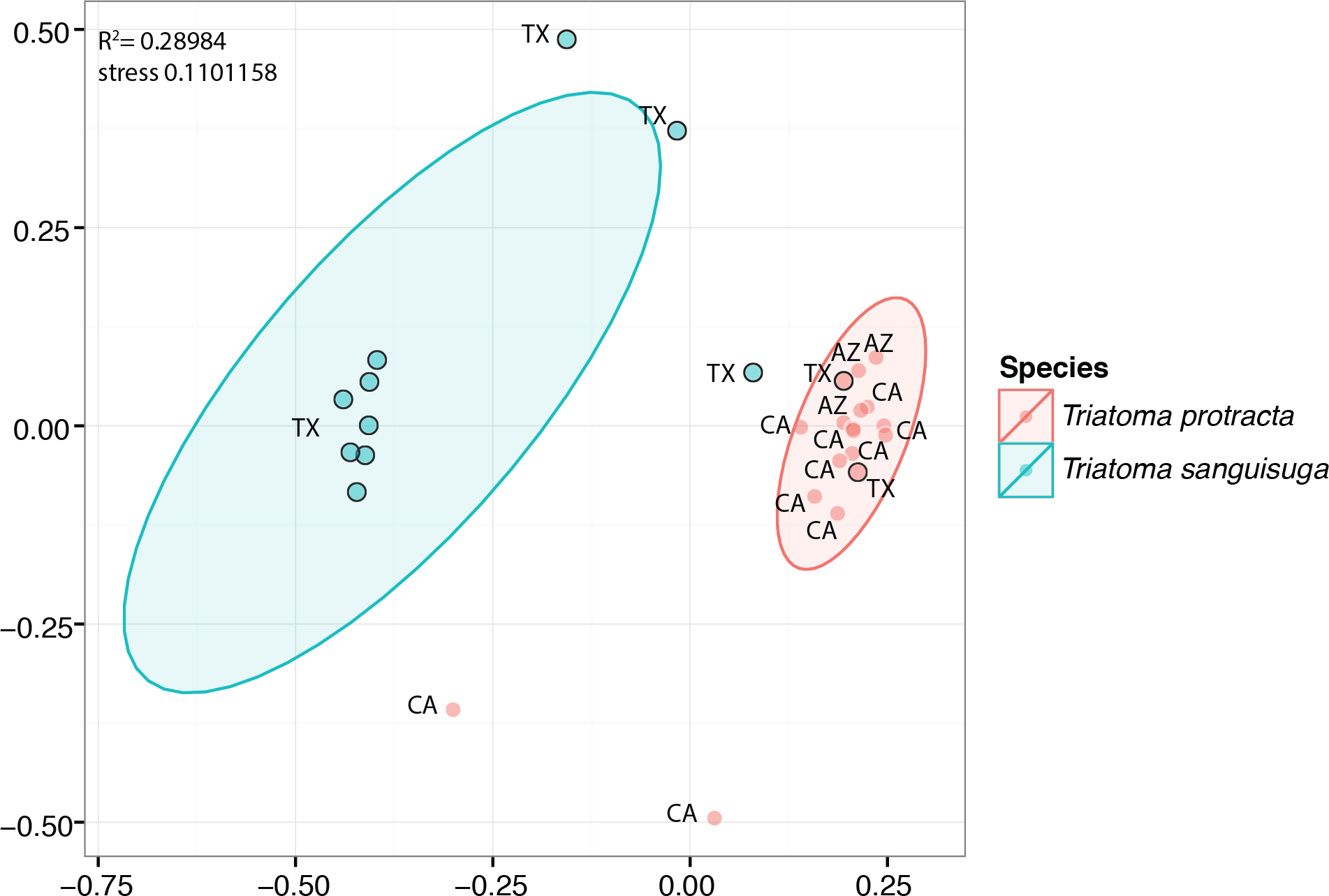
NMDS visualization of significant differences between the microbiomes of *T. protracta* and *T. sanguisuga* sampled from wild populations. The abbreviations stand for the states where the bugs were sampled. The black border circles designate samples for which the bioinformatic abdomen was generated (see Material and Methods section).

When testing possible variability of *T. protracta* microbiome composition in relation to the sex of the host individuals, their geographical origin or the year of sampling, no significant differences were found. The only significant correlation was identified for the presence of *T. cruzi* and the richness of *T. protracta* microbiome (Kruskal-Wallis chi-squared = 7.4558, df = 1, p-value = 0.006323; Figure 4). The prevalence of *T. cruzi* in 16 analyzed *T. protracta* adult individuals was 25%, with two cases of DTUI and the other two from DTUII-VI group. Due to a single case of *T. cruzi* infection in the tested set of *T. sanguisuga*, any correlation of microbiome profile with the presence of the parasite could not be evaluated for this species. It is however noteworthy that this case represented a double infection, with positive diagnostic PCR for both DTUI and DTUII-VI groups.

**Figure 4.**
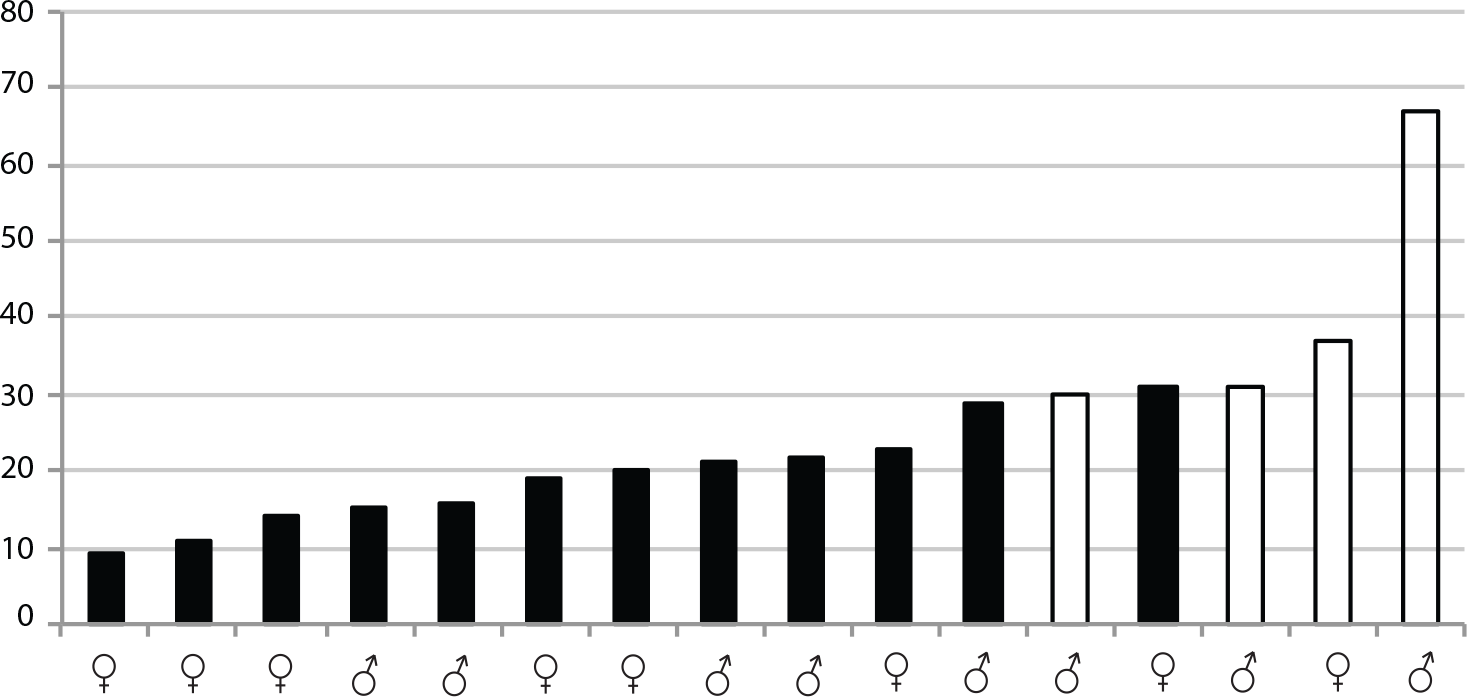
Significant differences in microbiome richness of *T. cruzi* infected and uninfected *T. protracta* adult individuals measured by Kruskal-Wallis rank test (chi-squared = 7.4558, df = 1, p-value = 0.006323). The numbers stand for the richness values. The black bars represent *T. cruzi* negative samples. The white bars are *T. cruzi* positive samples.

### Microbiomes in the laboratory reared colonies

The laboratory reared colonies also showed differential patterns in their microbiomes according to the Triatominae host species. Particularly, differences among the four species were evaluated as significant at the level of 0.001 (R^2^ = 0.784) using adonis (Figure 5). Colonies of *T. protracta* and *T. recurva* were dominated by Enterobacteriaceae (*Pectobacterium* sp.), while colonies of *T. vitticeps* were dominated mainly by Enterococcaceae (*Enterococcus* sp.) and also by Enterobacteriaceae (*Arsenophonus* sp.). *R. prolixus* did not show one single predominant OTU, but three with similar high abundance: Staphylococcaceae (*Macrococcus* sp.), Enterobacteriaceae (*Arsenophonus* sp.) and Moraxellaceae (*Acinetobacter* sp.). Bacillaceae (*Bacillus* sp.) were the next more predominant bacteria in *R. prolixus* and *T. vitticeps*.

**Figure 5.**
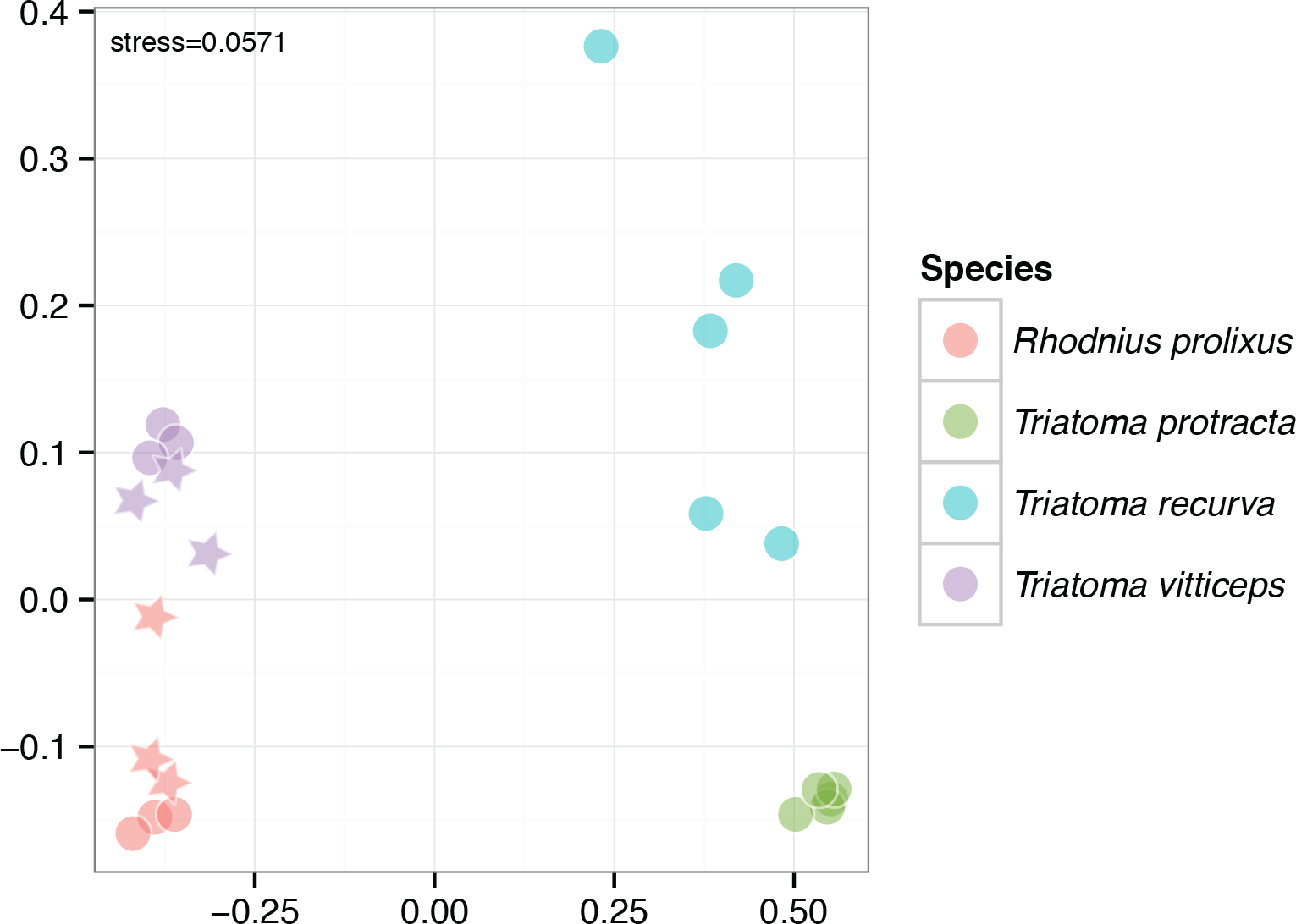
Significant differences among microbiomes of the lab-reared colonies depicted by the NMDS clustering (significance level of 0.001, R^2^=0.784). The circles stand for L1 instar, stars for L2.

### Arsenophonus triatominarum *patterns*

Both field collected and laboratory reared Triatominae showed distinct patterns for *A. triatominarum* presence. The averaged relative abundance of this bacterium in the gut of *T. sanguisuga* field-collected individuals was 39.76% (SD = 40.19%). However, no reads for *A. triatominarum* were detected in *T. protracta* from wild populations. For laboratory-reared colonies, the patterns of *A. triatominarum* were more variable: *R. prolixus* harbored *A. triatominarum* in the gut (26.36 ± 20.43%), but this bacterium was absent in the samples from eggs or faeces. *T. vitticeps* also harbored *Arsenophonus* in the gut (29.54 ± 28.21%). *Arsenophonus* was not found in *T. recurva* and *T. protracta* microbiomes of the sampled colonies.

## Discussion

### Microbiomes and host ontogeny

The microbiome is known to change along development in several arthropod species (Duguma et al., 2015; Gimonneau et al., 2014; Sudakaran et al., 2012; Zolnik et al., 2016). For example in ticks, the microbiome changes along the development towards higher diversity, probably due to off-host environmental bacterial acquisition (Zolnik et al., 2016). However, ticks feed on blood throughout their life, and the main component of the microbiome, *Rickettsia*, is present in all life stages, particularly in adult females and larvae, where it comprises most of the microbiome, possibly due to vertical transmission mechanisms (Zolnik et al., 2016). That is not the case in mosquitoes, whose larval microbiome is more diverse and highly influenced by the environment (i.e. aquatic), while the adult microbiome shows a different, more stable pattern (Duguma et al., 2015; Gimonneau et al., 2014). As mosquitoes are hematophagous only in the adult stage, it has been suggested that selection towards functionally important bacteria in this context acts to shape the adult female mosquito microbiome (Gimonneau et al., 2014). In the same way, hemipteran firebugs (*Pyrrhocoris apterus*) as a more phylogenetically (but not ecologically) related group to Triatominae, also
show differences when comparing the first instars with the latter instars and adults, probably due to the change of feeding habits between those stages (Sudakaran et al., 2012). The pattern we found in Triatominae, with the microbiome of early nymphs being different from the adults one, resembles that of the firebugs rather than that of the bloodfeeding insects (and more broadly arthropods). There is currently too little data to make any conclusion on the phylogeny or feeding strategy as the main determinant of the microbiome ontogeny, but it is interesting to note that even in strictly hematophagous vectors, the blood meal effect seems to be quite limited (Zolnik et al., 2016).

Triatominae can act as vectors of *T. cruzi* at any of their developmental stages. The prevalence of *T. cruzi* infection seems to be higher in adults and the latest instars compared to the younger nymphs, although this is based on a small number of unidentified nymph samples (Curtis-Robles et al., 2015; McPhatter et al., 2012; Wozniak et al., 2015). It is reasonable to suppose that these different infection rates are at least partially related to the different exposure to blood feeding on different hosts. Based on our analyses of the laboratory-reared individuals, we propose the changes of microbiome during the ontogeny as another possible determinant. A further investigation on the microbiome ontogenic patterns in wild populations of Triatominae and its correlation with *T. cruzi* infection is needed to confirm this hypothesis.

### Microbiomes in wild populations

Our results support previous findings on low-level complexity microbiomes of Triatominae. The most abundant bacterial taxa in *T. sanguisuga* and *T. protracta* comply with the diversity span previously described for South American species. *T. sanguisuga* microbiome profile dominated by two taxa, i.e. *Arsenophonus* and *Acinetobacter*, mirrors for instance that of *T. brasiliensis* complex (Díaz et al., 2016). *T. protracta* harbors distinct bacterial lineages. While *Salmonella* association with some Triatominae species was previously reported (Amino et al., 1998), additional taxa found here in *T. protracta* microbiome, i.e. Porphyromonadaceae (*Proteiniphilum* sp.), Clostridiales (*Peptoniphilus* sp.) and Neisseriaceae were identified as symbionts in other insects (e.g. Husseneder et al., 2017; Kwong et al., 2014; Zhang et al., 2017).

Microbiome species-specificity have been detected in several insect hosts to date (Aksoy et al., 2014; Leonhardt and Kaltenpoth, 2014; Novakova et al., 2017). The microbiome of Triatominae does not seem to be an exception, and species-specific patterns have been described as one of the main factors shaping the microbiome composition (da Mota et al., 2012; Díaz et al., 2016). While these studies were conducted mostly using laboratory-reared bugs, our results point out microbiome species-specific patterns in wild populations of NA 4 Triatominae.

Apart from the species-specific patterns, we also checked for other factors possibly affecting the microbiome composition in Triatominae. *T. protracta* was collected from three remote locations in the southern US (San Antonio, TX; Tucson, AZ; Escondido, CA). However, neither the geographical origin of the samples nor the sex of the adult individuals showed an effect on the Triatominae microbiome. These results are in accordance with the characteristics described for another hemipteran, the firebug, where the microbiome is remarkably stable across different populations, sexes, and even diets (Sudakaran et al., 2012).

### *Microbiome* T. cruzi *interface*

The prevalence of *T. cruzi* in our *T. protracta* samples was 25 %, lower than the prevalence found in previous studies of Triatominae from Texas (Curtis-Robles et al., 2017b). However, we still identified different *T. cruzi* DTUs (here only differentiated as DTUI and non-DTUI). These findings are in accordance with recent studies, where at least four different DTUs have been found in Southern US (Garcia et al., 2017). Particularly, the main DTU found in Texas (DTUI) bears a high medical importance as causative agent of Chagasic cardiac disease, highlighting the necessity for further research on this topic in the US (Curtis-Robles et al., 2017b; Garcia et al., 2017).

Previous studies have found a significant change in the microbiome composition of different South American species of Triatominae when challenged with a *T. cruzi* infected blood meal (Díaz et al., 2016). Working with wild populations, we collected *T. cruzi* positive and negative individuals, which allowed us to assess the correlation between the microbiome and the infection status of *T. protracta* adults. We found that the diversity of the microbiomes in *T. cruzi* positive individuals is significantly higher than in the negative ones. This result remarkably agrees with the findings on *T. cruzi* challenged *T. brasiliensis*, *T. sherlocki* and *P. megistus* (Díaz et al., 2016). However, due to the limited number of available samples, further studies will be necessary to confirm this correlation in natural Triatominae populations.

### Arsenophonus triatominarum

*A. triatominarum* has been described as an intracellular symbiont with strict specificity to various organs and tissues in *Triatoma infestans* (Hypša, 1993). It is therefore interesting to see this bacterium as one of the dominant components of Triatominae gut microbiomes. The presence of *A. triatominarum* in the host guts, and faeces, could provide an explanation for the intriguing host distribution of these bacteria. It has been noted that *A. triatominarum* is exclusively bound to Triatominae, but does not show phylogenetic concordance with the host (Šorfová et al., 2008). In fact, the extremely low molecular diversity of the Triatominae bound *Arsenophonus* suggests a recent acquisition and rapid horizontal spread rather than a long time coevolution. While the supposed intracellular location within internal organs was difficult to reconcile with such frequent and host specific horizontal transfers, the occurrence in gut content offers a new possible explanation, such as coprophagy or environmental contamination. Another interesting aspect of *A. triatominarum* being part of the gut microbiome is the possible role of this symbiont. While several members of the genus *Arsenophonus* are known for their profound effect on the host (reproduction distorters or nutritional mutualists; Wilkes et al., 2011; Nováková et al., 2015, 2016), no such phenomenon has so far been observed for *A. triatomiarum*. Considering a possible role of this symbiont, it might be interesting to note that unlike the other two major OTUs, *Enterococus* and *Acinetobacter*, its relative abundance seems to grow from the first until the third nymphal stages and then drops afterwards. The study of this symbiotic bacterium in the context of the complex and dynamic gut microbiome thus offers a new opportunity to search for its possible effects on the host fitness.

## Conclusion

The results presented here demonstrate that the composition of Triatominae microbiome is host-specific, maintaining its identity across large geographic areas. Compared to many other insect microbiomes, its overall diversity is low, with 12-17 main bacterial OTUs identified in wild populations of *Triatoma sanguisuga* and *T. protracta*. Surprisingly, one of the dominant taxa is the symbiotic bacterium *Arsenophonus triatominarum*, previously known as an exclusively intracellular symbiont from several Triatominae species. Based on the observations of a laboratory reared colony of *Rhodnius prolixus*, the frequency of the most abundant bacterial genera (*Enterococcus*, *Acinetobacter* and *Arsenophonus* in this Triatominae species) undergoes significant and consistent changes during the host ontogeny. Finally, a screening of *T. protracta* indicates possible relation between the microbiomes diversity and the occurrence of *Trypanosoma cruzi* in the host gut. This finding highlights the significance of further studies on the microbiomes of NA Triatominae species as a background for developing efficient vector control strategies.

## Acknowledgments

We would like to thank all the collaborators, listed in the manuscript, who provided us with the Triatominae samples analyzed in this study. The work was supported by the Ministry of Education, Youth and Sports of the Czech Republic, grant LH-15161 under the program for international collaboration KONTAKT II.

## Author Contributions Statement

EN and SR designed the study. RR, JS and WR managed the field collections and colony rearing. VS, SR and EN generated the data and performed the data analyses. EN, SR and VH drafted the manuscript. All the authors read and contributed to the final text.

